# LOGAN: A framework for LOssless Graph-based ANalysis of high throughput sequence data

**DOI:** 10.1101/175976

**Authors:** Anthony Bolger, Alisandra Denton, Marie Bolger, Björn Usadel

## Abstract

Recent massive growth in the production of sequencing data necessitates matching improvements in bioinformatics tools to effectively utilize it. Existing tools suffer from limitations in both scalability and applicability which are inherent to their underlying algorithms and data structures. We identify the key requirements for the ideal data structure for sequence analyses: it should be informationally lossless, locally updatable, and memory efficient; requirements which are not met by data structures underlying the major assembly strategies Overlap Layout Consensus and De Bruijn Graphs. We therefore propose a new data structure, the LOGAN graph, which is based on a memory efficient Sparse De Bruijn Graph with routing information. Innovations in storing routing information and careful implementation allow sequence datasets for *Escherichia coli* (4.6Mbp, 117x coverage), *Arabidopsis thaliana* (135Mbp, 17.5x coverage) and *Solanum pennellii* (1.2Gbp, 47x coverage) to be loaded into memory on a desktop computer in seconds, minutes, and hours respectively. Memory consumption is competitive with state of the art alternatives, while losslessly representing the reads in an indexed and updatable form. Both Second and Third Generation Sequencing reads are supported. Thus, the LOGAN graph is positioned to be the backbone for major breakthroughs in sequence analysis such as integrated hybrid assembly, assembly of exceptionally large and repetitive genomes, as well as assembly and representation of pan-genomes.

## Introduction

DNA sequencing is currently undergoing a second revolution. The first brought massively parallel short-read sequencing platforms, now called Second Generation Sequencing (2GS). The current revolution, Third Generation Sequencing (3GS), operates on single molecules and produces much longer reads, on the order of 10s of kilobases, but at a much higher error rate. These new sequencing technologies produce massively more data for less time and cost, shifting the bottleneck from wet-lab work towards the bioinformatic steps that follow. As a result, there is an urgent need to develop tools that can (i) efficiently handle large sequencing datasets, (ii) utilize the complementary nature of 2GS and 3GS data, and (iii) ultimately deliver improved results with less human and computational resources.

## First Principles Analysis

The analysis of sequence datasets is, in principle, the distillation of a very large number of low information, error-prone pieces of evidence (e.g. reads) into a modest number of high information, highly reliable statements (e.g. long contig sequences). We concur with Medvedev and Brudno ^(1)^ that the goal should be the set of contigs which best explains the reads, given reasonable priors regarding their creation, thus aiming towards a parsimonious result, rather than one of strictly minimal length. The information in reads originating from overlapping regions in the origin (e.g. genome) sequence can be combined to produce high confidence contigs provided three key criteria are met. These are that a) a series of overlapping reads exist spanning the origin sequence, b) true overlaps and false overlaps can be differentiated, and c) the overlaps can be used to correct errors in the reads.

The various sequencing platforms produce read datasets of widely varying characteristics, including the dataset size and cost, but from an analysis perspective, the key attributes are the read length and error rate. Current datasets can consist of short, relatively accurate reads from 2GS sequencing platforms and/or longer, more error-prone 3GS reads.

One major confounding factor is the presence of repetitive regions in the origin sequence. Unambiguous reconstruction of such regions, which consist of multiple (identical or near-identical) copies of a subsequence, requires reads of sufficient length to span the identical regions. Thus the severity of the problem posed by repeats is a function of the relative length of repeats and reads. A second potential confounding factor is the relative abundance of origin molecules, i.e. the dynamic range, which complicates differentiating sequencing error from rare sequence variants. This is of particular relevance in the analysis of transcriptomic or meta-genomic datasets.

Combining the information from multiple reads requires gathering similar sequences, and evaluating whether differences are due to sequencing error or true variation. In the case of sequencing error, the variation is incorrect and should be removed, while in the case of true variation, the reads should be partitioned and used to reconstruct their distinct origins independently. While accurate analysis relies on distinguishing read error from true variation, solving either aspect is non-trivial without correct prior resolution of the other. This inherently circular nature suggests an iterative, incremental approach, interleaving partitioning and correction, rather than performing them sequentially.

## Computational Considerations

The essential challenge, from a computational perspective, is one of scale. The core operations required, such as sequence comparison and alignment, already have well-established algorithms that guarantee optimal solutions ^(2,3)^. However, these approaches are not computationally practical for non-trivial datasets. Therefore, a plethora of specialized alternative approaches have been developed, which offer massive computational benefits in time and/or memory, typically in exchange for a moderately sub-optimal result. To better explain the innovations proposed in this paper, the most relevant existing assembly approaches will be briefly described. A full dissection of such approaches is beyond the scope of this paper, but can be found in comprehensive reviews such as Miller et al. ^(4)^ and Koren and Phillippy ^(5)^.

## State of the Art

### Overlap-Layout-Consensus

The traditional approach to *de novo* assembly is Overlap-Layout-Consensus (OLC), named after the three main analysis processes involved. In the overlap phase, reads are compared pairwise, to determine if they potentially came from overlapping locations in the origin. The output of the overlap phase is a graph where each read is represented as a node, and each detected overlap is an edge. The layout phase uses the overlap graph to partition the read dataset and create a putative layout of the reads. Finally, the consensus stage resolves any disagreements between the read of overlapping origin, which are presumably caused by sequencing errors, and thus produce an accurate final sequence.

The OLC approach is relatively tolerant of errors in the reads. However scaling OLC to the large datasets from whole-genome shotgun sequencing of highly repetitive targets is challenging. As a result, this approach had fallen out of favour during the 2GS era, but has regained popularity since it is more suited to the high error rate and long read characteristics of 3GS datasets.

### Detailed Assessment

OLC uses the ‘partition before correction’ strategy, which has both superficial and fundamental problems with repetitive regions. One superficial problem is the storage requirements for the overlap graph, which scales quadratically, i.e. *O*(*N* ^2^) with coverage N, and in effect, the coverage of each repeat region is multiplied by its copy number. This can be reduced to linear scaling, i.e. *O*(*N*) with coverage N, by transitive reduction, proposed in Myers ^(6)^, and used in the Burrows-Wheeler Transform ^(7)^ (BWT) based String Graph Assembler ^(8)^. Another superficial problem is that optimizations to speed up the pairwise overlap step, such as finding shared seeds, suffer from high false positives in repetitive regions. This can be reduced by filtering or down-weighting commonly found seeds ^(9)^.

The fundamental problem is that accurate partitioning of the error-containing reads into subsets with a single origin is not always possible. False negatives caused by read errors may result in missing valid overlaps. While reads may be corrected earlier, to do so validly requires an approximate partitioning, followed by conservative error removal. This is in effect a preliminary application of the OLC approach to produce reads with fewer errors, and may be repeated to further correct the reads. On the other hand, reads from very similar repeat copies will be effectively indistinguishable, and result in false positives. Thus sequences from multiple target regions will be included in a single layout and consensus task. To avoid the creation of a collapsed repeat, the layout and consensus steps must detect divergent sequences, and group them appropriately, in effect, using a nested OLC-like step. Thus the circular nature of the problem causes an apparently single pass partition-first approach to become an iterative (and inefficient) partition and correction process.

### De Bruijn Graph

Another well-established method for *de novo* assembly is the De Bruijn Graph (DBG) approach, named after the intermediate data structure used to represent the sequence information. A DBG is constructed by breaking each read into a series of overlapping *k*-long subsequence fragments, known as *k*-mers. The graph is then populated with *k*-mers as nodes, and edges connecting the nodes from successive *k*-mers. Nodes and edges which occur in multiple reads are re-used, and thus the memory requirements are independent of coverage, i.e. *O*(1), at least for error-free sequences.

Unlike the OLC approach, the DBG approach scales well with read number and therefore became dominant during the 2GS sequencing era; although the earliest implementations, such as the Euler assembler ^(10)^, pre-date the availability of 2GS platforms.

### Detailed Assessment

The DBG approach uses an incremental multi-stage analysis strategy. A preliminary partitioning occurs implicitly during graph creation, with shared *k*-mers from different reads merging into the same nodes and edges. Erroneous nodes and edges are detected using various rules and heuristics, often based on coverage and/or graph topology. Linear paths through the remaining nodes and edges are determined, and extracted as contig sequences. Additional information, such as the original read sequences or read pairing can be used to guide the contig extraction process through otherwise ambiguous regions.

The DBG approach also has both superficial and fundamental problems. One superficial problem is the memory consumption of the graph, since it stores the primary sequence data in memory, and it contains, informationally speaking, a high level of redundancy. In the purely theoretical case of error-free data, the resulting expansion would be independent of coverage and bounded as *k* times the target size. However, in practice reads are not error free, and a single base error generally causes the formation of *k* erroneous *k*-mers, and furthermore, this effect scales linearly with coverage. As such, the number of error-related nodes can be estimated as the product of *k*, coverage and error rate. Therefore, even with relatively low error rates, in the order of 0.1-1%, assuming typical *k* value of at least 31, and coverage levels of ~100x, error-related nodes make up the vast majority of the DBG. Newer implementations have reduced this memory requirement substantially, although at the cost of computational overhead and/or early data filtering.

The fundamental problem is that two important aspects of the graph are determined by a single user-chosen *k* parameter. The first aspect is the minimum length of identical sequence required to merge reads into a shared node, which is equal to *k* itself. The second aspect is the maximum length of sequence retained within the graph, generally *k* + 1, corresponding to an edge. Tolerable tradeoff values for 2GS datasets are generally possible, since the reads are relatively accurate, so merge sensitivity is not critical; and the reads are relatively short, limiting the amount of information lost due to the fragmented representation. For 3GS datasets however, there is greater need for the former to be relatively small, to allow for merging despite higher error rates, while one would like the latter to be large, conserving more of the information within the long reads. The imposed linkage between these properties makes the DBG entirely unsuited to 3GS data.

This information loss, which is inherent in the DBG, has a critical effect on the analysis. As a simple example, consider a case where two independent reads form a ‘frayed rope’ structure in the graph, as illustrated in Figure 1. Each read has independent nodes at its start and end, but shares one or more nodes in the middle. From the DBG alone, it is not possible to distinguish the paths of the original reads (Interpretation 1), from the ‘spliced’ possibilities or phantom paths, which mix the pre-repeat and post-repeat sections of different reads (Interpretation 2). Thus any analysis based on the DBG alone is limited to incomplete information. Although this effect can be reduced by using a larger *k* value, other practical factors such as error rate and coverage typically limit the maximum *k* value to substantially shorter than the read length. Given the drastic consequences of this information loss, many assemblers attempt to re-map the original reads onto the graph later in the analysis. However, this can only reduce, not fully eliminate, the negative effects.

**Figure 1:**
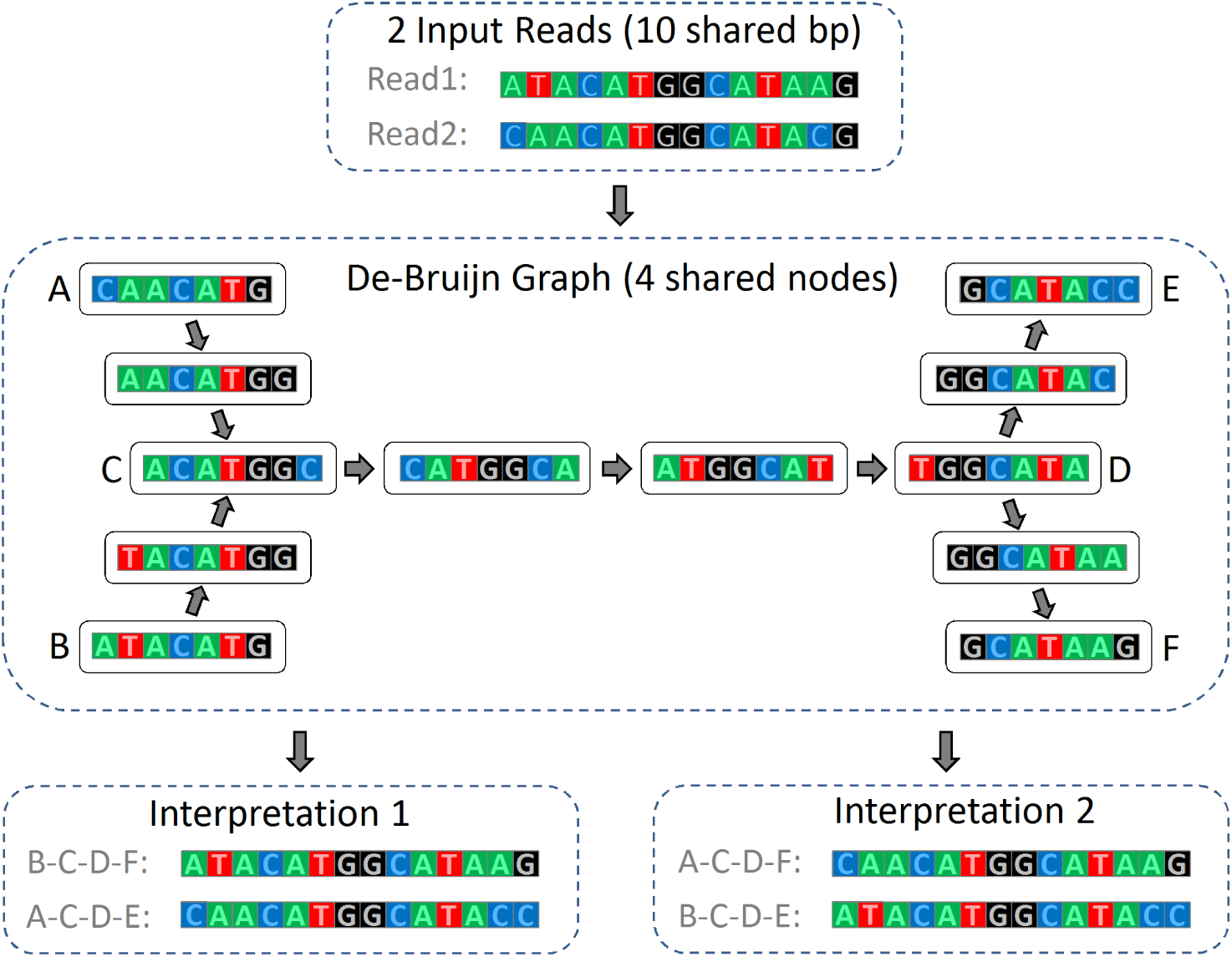
An example of a ‘Frayed Rope’ in a De Bruijn Graph: The two reads differ on each end, but share a 10 base subsequence. This length is equal to *k+3*, and thus the reads share 4 nodes. The resulting graph is compatible with two interpretations, with the first, on the left, being correct. Interpretation 2, on the right, in incorrect since it follows ‘phantom paths’ which splice the reads.

### Space-Efficient DBG Representation

The memory consumption of naïve implementations of the DBG has motivated space-optimization of the data structure. One approach is to represent the nodes or edges of the graph more efficiently. Conway’s ^(11)^ representation uses a Succinct ^(12)^ approach to store the edges, in both original and reverse-complemented forms. In contrast, Minia ^(13)^ stores only the nodes, using a cascading Bloom filter approach.

Another approach, the Sparse graph ^(14)^, exploits the redundancy between successive *k*-mers, by creating nodes only for a subset of the *k*-mers. Although this results in more information in the edges, the Sparse graph has less redundancy than the standard ‘dense’ version. The sparse approach introduces a second configurable parameter, *g*, indicating the maximum number of bases between successive *k*-mers, i.e. the maximum edge length. The restriction *g < k* is proposed, but is not conceptually essential to the approach. If *g* = 1, the sparse graph is equivalent to the standard DBG. Note that the sparse graph is not, in general, equivalent to the DBG, since it cannot efficiently answer presence/absence queries on all *k*-mers in the dataset.

### Towards a Grand Unified Approach

As previously described, sequence analysis is inherently a circular problem, and cannot be optimally solved with an a-priori ordered list of operations. The alternative is an iterative progressive refinement approach, which opportunistically improves the solution during each iteration. The most apparent errors corrected and the most similar sequences can be identified and handled during the initial rounds of improvement. In turn, these modifications enable further changes, until the solution converges, hopefully providing a high quality and accurate result.

Progressive refinement can be efficiently implemented using a data-centric architecture, such as the Blackboard approach (used by the Hearsay-II speech recognition system ^(15)^) and other similar software design patterns ^(16)^. These place the representation of the dataset under refinement at the centre of the solution, with multiple independent ‘actors’ concurrently accessing and updating the shared data representation. These ‘actors’ combine in an ad-hoc manner to progressively refine the solution. Example modifications include changing a sequence caused by an error, grouping reads with putative common origins and identifying putative longer-distance arrangements. The structure of the dataset is conserved throughout the process, allowing ad-hoc ordering between transformations. This data-centric approach can be contrasted with the more typical process-centric pipeline approach, which applies a directed series of transformations, and often changes the structure of the data representation during or between transformations.

A fundamental roadblock is a suitable representation of the dataset during this process. It must provide ‘mechanism without policy’, in that it should not limit information available to the ‘actors’ nor the type of transformations that can be applied, when compared to performing the same transformations in a serial pipeline. This implies that the representation be informationally lossless, with the full initial sequence dataset available for analysis. A second condition is that the representation support appropriate update operations, allowing inter-actor visibility of updates.

Furthermore, the representation must be computationally reasonable, in terms of both processor and memory requirements. Ensuring processor efficiency requires that the data representation be locally updatable, such that each modification affects a small portion of the dataset. Memory requirements are primarily driven by the need to support random access patterns within the entire dataset, and thus must be stored in primary memory (RAM). Given the size of raw sequence data alone typically exceeds most computers’ memory capacity, and since substantial auxiliary data is needed during analysis (generally much larger than the raw sequences), the required representational efficiency is extremely challenging to achieve.

We can therefore, identify the following core requirements:

- Lossless: It must be capable of representing input sequences of any length without modification or data loss.
- Efficient: Datasets must be stored in an space-efficient manner, allowing very large datasets to be analysed using modest computational resources.
- Locally updatable: Modification of sequences, as a result of e.g. error correction, should require limited changes to the data representation.

The OLC approach is process centric, and therefore contrary to the progressive refinement approach. In contrast, the DBG approach is data-centric and offers the possibility of computationally efficient, iterative refinement of sequence data in an in-memory structure. However, its inherent and drastic information loss nullify this advantage. Furthermore, as previously noted, it is also an informationally inefficient representation, typically requiring much more memory than the raw sequence data.

Here we present the LOGAN graph, which extends the DBG approach to meet the above requirements, providing a data structure suitable for a wide range of data-centric sequence analysis applications.

## Materials and Methods

### Extending the DBG

The DBG approach offers one of the three requirements, local updatability, but lacks the informational lossless property, and struggles with memory efficiency. This is related to two distinct effects of the *k* parameter previously noted. Low *k* increases merge sensitivity, thus improves error tolerance, and also reduces memory requirements (particularly considering errors), both of which are desirable. On the other hand, a high *k* retains longer-distance information, which is also desirable. This is because only the set of nodes and/or edges traversed by the combined dataset are recorded in the graph topology, while the specific paths taken by each individual read is generally lost, although it can be inferred in some cases.

Retaining the paths taken by individual reads through shared nodes/edges would make the DBG a lossless datastructure. We refer to this as ‘routing’, since it is analogous to routing in communication networks. The routed DBG thus meets the informationally lossless requirement. Furthermore, alleviating information loss is the only motivation for using larger *k* sizes. By sidestepping this problem entirely, a much smaller *k* size can be used, reducing the storage costs of *k*-mers, especially in the case of errors, helping meet the third requirement.

### Route Representation

Since the DBG is generally stored in memory, and represents the entire dataset, space-efficiency is a key concern. Furthermore, unlike the nodes and edges which are shared between multiple sequences, routing information is required for each input sequence. As a result, efficiently representing the routing information within a DBG (or DBG-like graph) is absolutely critical. The previously shown ‘frayed rope’ problem, and some minor variations thereof, will be used to illustrate various methods of storing the routing information, since the general form of this problem, with an arbitrarily large number of shared nodes or sequences, is sufficient to assess any proposed solution.

Given such a ‘frayed rope’ structure, one possibility is to enumerate the set of edge-pairs which correspond to valid transitions across a given node, as shown in Figure 2. Unfortunately this resolves only the ‘single shared node’ case of the ‘frayed rope’, shown on the left. If the reads converge in one node, and diverge in another node, it is not possible to uniquely resolve the original reads, as shown on the right for the ‘two shared node’ scenario. Each edge-pair in the routing table in fact represents a sequence of *k* + 2 bases, an improvement on the standard DBG, where edges represent *k* + 1 bases, but hardly dramatic. Extending this approach to store additional context from further ‘upstream’ merely incrementally improves the length of shared sequence which can be resolved, at the cost of intolerable duplication between the routing tables of the various nodes.

**Figure 2:**
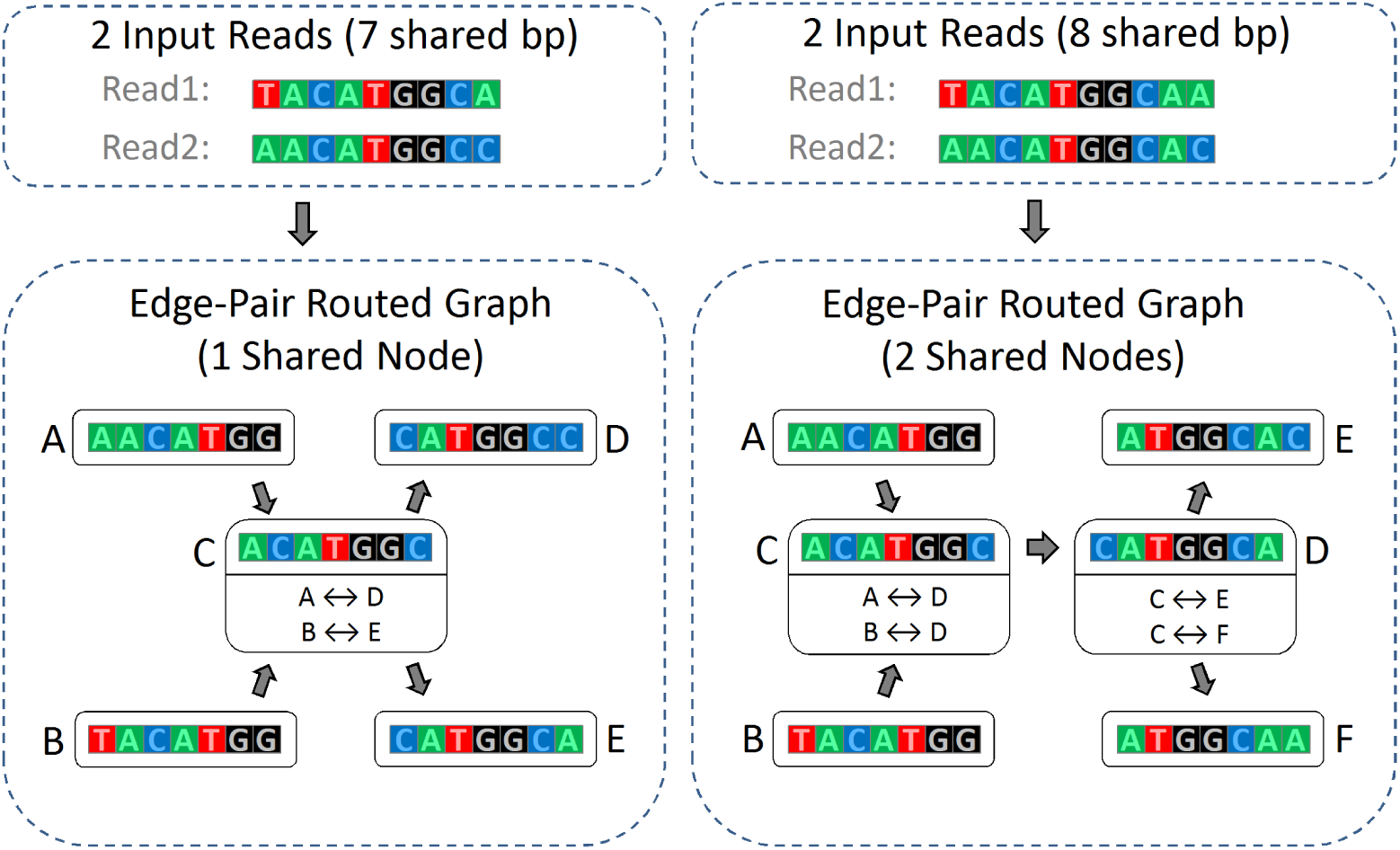
Edge Pair Routing Example: The left side shows two reads which differ on each end, but share a 7 base subsequence. This length is equal to *k*, thus the reads share one node. The resulting graph can be disambiguated using the edge-pair routing table shown in Node C. The right side shows two reads which share a 8 base subsequence. This length is equal to *k+1*, thus the reads share two nodes. The edge-pair approach is unable to resolve this scenario, since there is no defined association between the edge-pairs in node C and those in node D.

The problem of determining the validity of long-distance paths can be resolved by associating the edge-pairs stored in different nodes which relate to a single read. One approach would be to use a unique global identifier for each read, and thereby associate each edge-pair with the reads from which they originate. Thus when walking the graph, the edge-pairs from each read can be associated and used to determine which successor nodes to choose at each step, and thus longer distance paths which lack any read support can be avoided. This approach however would massively increase the memory requirements, since each node transition of every read would require an additional read identifier in the graph. Since the number of reads in the graph is typically large, and the relationship between read identifiers and nodes is effectively random, the space requirements for these identifiers would be prohibitive.

A third alternative, which was chosen here, is to use local read identifiers, assigned independently in each edge. This approach is analogous to label-switching network technologies, such as ATM (Asynchronous Transfer Mode) ^(17)^ or MPLS (Multiprotocol Label Switching) ^(18)^. Each node contains a routing table, with each route mapping between a left-side edge-label pair and a right-side edge-label pair, shown as the upper (blue) routing table variant in Figure 3. Transitioning a node involves looking up the current sequence information in the routing table, based on the inbound edge and label, and determining the appropriate outbound edge-label pair, exactly analogous to routing process in label-switching networks. Since each edge-label pair occurs exactly once within a routing table, this approach fully resolves the ambiguity which occurs when a node with multiple outbound edges is encountered in a standard DBG. In storage terms, this approach is still somewhat inefficient, but since the labels are assigned on a per-edge basis, they are relatively small in the majority of edges.

**Figure 3:**
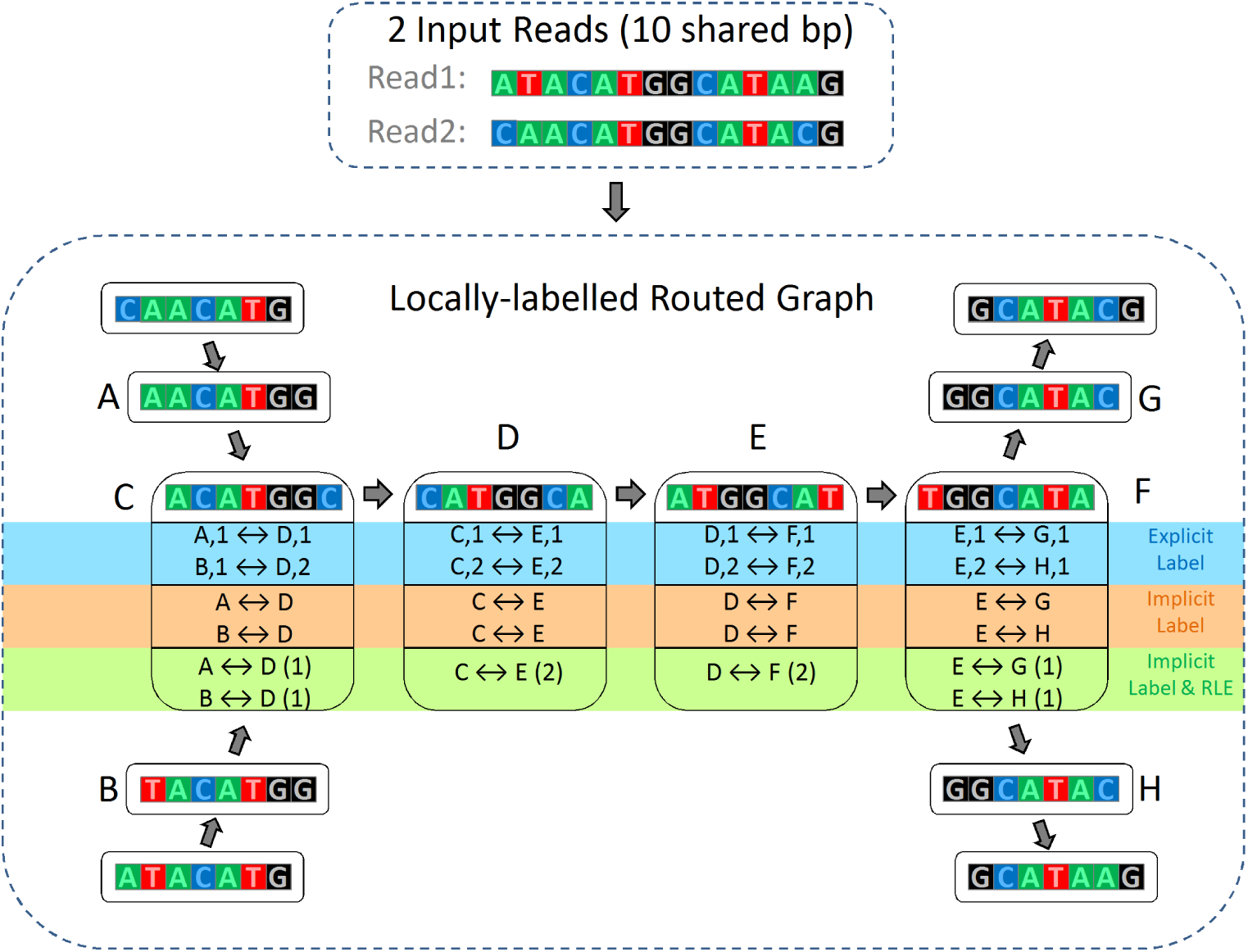
Local Label Routing Example: These reads share a 10 base subsequence, which is equal to *k+3*, and thus the reads share 4 nodes. The graph shows three variants of local label routing in Nodes C-F, with the trivial routing information from the other nodes omitted for brevity. The upper routing table in each node, highlighted in blue, shows the ‘explicit label’ approach, which assigns a specified local label to each sequence. The middle, highlighted in orange, is the equivalent but more efficient ‘implicit label’ approach. The local label is determined implicitly based on the position of the route within the routing table. The lower routing table combines the implicit label approach with Run Length Encoding (RLE), with the count indicator in parenthesis. Non-trivial routing first occurs in Node C, where the numeric label 1 or 2 is assigned, depending on the prior node. This label is then used to transition the following nodes, to lookup the corresponding route entry and new label. In this example, the numeric label remains unchanged until node F, where it is used to determine if the outbound transition should be to node G or H.

### Further Refinements

A substantial efficiency improvement was achieved by considering that the label itself is irrelevant, provided that the same label for each sequence is used by both nodes at the end of a given edge. If route entries were always appended to the routing table, an implicit label could be assigned to each sequence, by simply counting the number of sequences which used the same edge already in the routing table (these will agree since the two routing tables must contain corresponding routes for all existing sequences spanning the edge). This implicit label can be left un-stored, and recreated on demand simply by counting the uses of the same edge earlier in each routing table. This approach is shown in the middle (orange) routing table variant in Figure 3.

For nodes with one or relatively few spanning reads, whether from errors or low-copy number sequences, this implicit label representation is reasonably space efficient. However, with a typical genome dataset, having a coverage of e.g. ~ 100*x*, the nodes which represent single-copy regions will contain many routes, each corresponding to the same left-side/right-side edge pairing. This will result in ~ 100*x* identical routes, with a small number of other routes, representing reads which diverge due to nearby errors. While little can be done about the error-related routes, the informational redundancy between identical routes was exploited using the run-length encoding ^(19)^ (RLE) approach, which replaces a series of consecutive identical entries with a single value and a count indicator. The combination of implicit labelling and RLE encoded route entries is shown as the lower (green) routing table variant in Figure 3, with the count indicator in parenthesis.

The append-only requirement proposed above is actually overly restrictive. The range of compatible positions in each route is reasonably flexible, and requires only that the new entry is placed consistently, within both routing tables, relative to existing entries referring to the same edge. It can be inserted earlier than or after all existing routes, or if placed between existing entries implicitly indexed as *X* and *X* + 1 in one routing table, it must be placed between the corresponding entries in the other routing table. This criterion is logically derived from the implicit labelling (number of routes using this edge earlier in the routing table) and the need for both nodes to agree on the label for each sequence within the edge. Naturally, inserting a new sequence before an existing sequence(s) assumes the label from that sequence, and causes the implicit labels of all existing later edges to be incremented. The presence of routes referring to different edges is irrelevant, since labels are edge-related.

In repetitive nodes, there may be several highly used edge pairings, each representing valid spanning of the node by reads originating in different repeat copies.These highly used pairings are accompanied by rarely used edge pairings, representing reads with nearby errors. Each of the commonly occurring pairings is a prime candidate for RLE, but achieving maximum RLE efficiency requires that the identical entries are grouped in the routing table. Since entries referring to independent edges can be ordered at will, this can generally be achieved by ordering the pairings based on their edges. As a result, the number of non-error entries in each node can be drastically reduced, and becomes almost independent of coverage.

## LOGAN Graph

The LOGAN Graph proposed here integrates a highly efficient representation of sequence routes into a space optimized DBG-like implementation, to create a practical lossless sequence representation and analysis structure. It is based on a ‘nodes plus embedded edges’ sparse graph representation. The term *k*-mer is used to refer to all *k* long sub-sequences within the dataset, while the term s-mer refers to the subset of *k*-mers which become nodes. Forward and reverse-complementary nodes are merged, with the canonical form chosen based on whether the forward or reverse packed form is numerically lower (equivalent to lexicographically lower, when written in text form).

Edges are stored per node, grouped into prefix and suffix edges. Since each node may be transitioned either canonically or non-canonically, depending on what is contained in the original sequence, a node can be transitioned prefix-edge to suffix-edge, or suffix-edge to prefix-edge. Therefore the terms ‘upstream’ and ‘downstream’ are used to indicate the edges from perspective of the spanning sequence. Edges may optionally lead to other nodes, via the canonical or non-canonical form, or may be dangling, indicating the end of the sequence.

Routing information is stored per-node, using the RLE-compressed implicit local label approach described above. Each node contains two routing tables, with sequences spanning the node in the canonical orientation stored in the ‘forward’ routing table, and those spanning in the non-canonical orientation in the ‘reverse’ routing table. The routing tables are sorted recursively based on the ‘upstream’ sequence path, with the ‘downstream’ path used as a tie-breaker.

Sequence ends are indicated using two methods. One is the dangling edge scenario, which is used if the final *k*-mer in the sequence is not a node. Alternatively the end can be indicated by a routing table entry referencing the ‘null’ edge, currently stored using edge index 0.

### Preliminary Assessment

The LOGAN graph is broadly based on the DBG, and aims to retain the benefits of that approach while overcoming the shortcomings. The most important benefits of the DBG approach are that it entirely avoids the *O*(*N* ^2^) processing and memory scalability issues of the OLC approach, and that is allows a data-centric analysis approach. These advantages are retained in the LOGAN graph.

The critical shortcoming of the DBG approach is the inherent information loss, with memory consumption an important secondary concern. Resolving these shortcomings requires a multi-faceted approach. Addition of routing data overcomes the fundamental issue of information loss. The biggest computational weakness, high memory consumption, is addressed by two factors. The first is the Sparse graph representation, which limits the number of new nodes created by a single base error, from *∼ k* in a DBG, which is typically at least 31, to typically 2 or less (depending on the sparseness allowed). The second factor is that *k* in a LOGAN graph can be much smaller, since reducing *k* does not cause information loss, due to the routing information. The smaller *k* means that substantially fewer nodes are created, since fewer unique *k* long sub-sequences exist in the dataset, and additionally the nodes themselves are smaller.

One rather theoretical advantage of the DBG, the coverage-independent storage requirement given error free data, is diluted slightly in the LOGAN graph. This is because, unlike nodes and edges which can be shared between input sequences at no additional cost, there is an overhead for extra sequences even if following identical routes. However, the use of RLE means that doubling the number of routes requires just one additional bit per route entry. Thus, in practical datasets, which are not error free, the cost of these wider route entries is expected to be trivial.

## LOGAN Graph Evaluation

To evaluate the LOGAN graph, we chose several public sequence datasets which covered a broad spectrum of species as test cases. These datasets consisted primarily of 2GS data from various Illumina platforms, but some tests used 3GS or hybrid datasets. Each dataset was then used to build a LOGAN graph representation, to assess graph build time and size of the resulting graph.

### Dataset Preparation

The 2GS datasets were trimmed using Trimmomatic ^(20)^, with appropriate trimming for TruSeq2 (ILLUMINACLIP:TruSeq2-PE.fa:2:30:12), TruSeq3 (ILLUMINACLIP:TruSeq3-PE.fa:2:30:12) or Nextera adapters (ILLUMINACLIP:NexteraPE-PE.fa:2:30:12), depending on the library preparation. Quality filtering was performed using end trimming (LEADING:3 TRAILING:3) and sliding window quality filtering to Q20 (SLIDINGWINDOW:4:20). Short reads and reads containing N bases were removed using additional filters (MINLEN:40 and BASECOUNT:N:0:0).

The 3GS dataset was filtered to remove reads shorter than 5kbp, but was otherwise used as is.

### Datasets

Public genomic datasets, covering a range of genomes, from the relatively small and simple *Escherichia coli* str. K12 genome (~ 4.6Mbp), through *Arabidopsis thaliana* cv. Col-0 (~ 135Mbp) to the more complex *Solanum pennellii* cv. LA716 (~ 1.2Gbp) were used to evaluate the LOGAN approach. The details of these datasets are given in Table 1.

**Table 1.** Evaluation Datasets: The size of the datasets, in millions of reads and megabases, plus the approximate genome coverage is shown for each dataset, before and after filtering. 2GS datasets were trimmed using appropriate adapter sequences and quality filtered using Sliding Window trimming at the specified threshold, with short (<40bp) or N-containing reads dropped. 3GS datasets were filtered for the specified length only.

| Dataset | Platform | Source | Filtering | Reads (m) | Bases (Mbp) | Coverage |
| --- | --- | --- | --- | --- | --- | --- |
| *E. coli* 1 | Illumina | SRX016044 | Unprocessed | 20.7 | 2,091.4 | 450.77 |
|  |  |  | Q20 | 7.6 | 543.7 | 117.19 |
| *E. coli* 2 | Illumina | SRX131029,<br>-33,-38,-53 | Unprocessed | 283.8 | 28,660.4 | 6177.25 |
|  |  |  | Q20 | 221.0 | 19,743.8 | 4,255.42 |
| *E. coli*<br>PB1 | PacBio | SRX255228 | Unprocessed | 1.39 | 2,611.6 | 562.9 |
| | | | Len ≥ 5000 | 0.15 | 1,060.5 | 228.6 |
| *A. thaliana* | Illumina | SRX015812 | Unprocessed | 34.7 | 3,468.4 | 25.69 |
|  |  |  | Q20 | 30.4 | 2,364.5 | 17.51 |
| *S. pennellii* | Illumina | ERR538235-40 | Unprocessed | 521.9 | 78,809.7 | 65.24 |
|  |  |  | Q20 | 450.3 | 56,658.8 | 46.90 |

## Results

### Prototype Implementation

Creating the LOGAN graph from a given dataset is theoretically simple, but in practice, a major computational challenge. Nonetheless, a reasonably efficient implementation has been developed, and will be described in detail in future. It is used here to evaluate the practicality of the LOGAN graph as a foundation for sequence analysis tools.

### Overview

As previously described, the LOGAN graph uses the canonical node form of the Sparse De Bruijn Graph, with edges embedded within nodes, extended by adding routing information to each node. Both nodes and edges are stored using a 2 bit per base packed representation. The current implementation does not exploit similarity between edges within a node, although this in principle could further improve representational efficiency. As previously described, forward and reverse routing tables are represented separately, and stored using variable width bit-packed representation. The number of bits allocated to each field (prefix edge, suffix edge and route entry width) is based on their corresponding maximum values in each routing table.

Both node-size *k* and maximum sparseness *s* are set to 23 in the current implementation. We use *s* rather than the *g* used in Ye et al. ^(14)^, to refer to maximum sparseness, for historical reasons. These values were chosen primarily for 2GS datasets, and it is anticipated that they could be altered in future, with larger *s* and perhaps a smaller *k* used for more efficient representation of 3GS and hybrid datasets, due to the higher error content.

The prototype uses a two-phase approach. The first phase is indexing, where a subset of *k* - mers within the dataset are selected to become s-mers. This is followed by routing, which involves adding each sequence to the spanned s-mers, creating edges and route entries as needed. Although it is in principle possible to perform these steps within a single pass, it is in practice somewhat easier to use a two-phase approach, where the entire dataset is indexed before routing is started.

### Indexing

The primary goal of indexing step is to determine a subset of *k*-mers to index as s-mers. The only requirement is that the maximum distance between successive s-mers in the dataset remains within the defined *s* value. Secondary goals include optimizing overall memory efficiency, by creating as few s-mers as possible, and by preferring s-mers which connect otherwise disconnected sequences, e.g. which share relatively short sequences, thus increasing merge sensitivity.

The current implementation uses a simple approach, as follows: For each sequence, all *k* - mers are determined and checked if they are already indexed. If a long stretch of non-indexed *k*-mers are found, the stretch is divided by indexing additional *k*-mers, such that the maximum distance between indexed *k*-mers is respected. A multi-threaded implementation can be achieved by processing independent sequence batches in each thread, while using a shared concurrent data structure to represent the set of already chosen indexed *k*-mers. However, this approach is non-deterministic, i.e. there is no guarantee that the same set of *k*-mers will be chosen on successive runs, given the same data.

### Routing

The primary goal of the routing step is to add each sequence in the dataset to the graph precisely, with the secondary goal of optimizing memory usage by arranging reads to fully exploit the compression potential of the RLE route entry representation. As previously mentioned, this is achieved by sorting all sequences within each routing table by their upstream sequences, ensuring groups of sequences which have been coherent previously, and are thus likely remain coherent in this node, can be efficiently represented by a single or at worst a modest number of routing table entries.

Sequences are inserted in their original orientation i.e. 5’ to 3’, with the goal of achieving their preferred relative upstream ordering, with the downstream sequence used as a tie-breaker. An apparent problem is that the relative placement of sequences cannot always be fully resolved locally, i.e. within a single node, since the sequence deviation which determines their ultimate ordering may not occur within this node. Scanning ahead to determine the ordering would be possible but highly inefficient.

The solution to this issue is rather subtle: the sequences can be added to the graph without determining their exact relative position, since the update to initial node(s) will be sufficiently ambiguous to represent any relative ordering which is determined later. Therefore, during the insertion process, the edge position of the new sequence is stored not as a single value, but as a range. This range generally narrows progressively as the existing sequences diverge, since these divergences invalidate some of the possible positions (since such positions would no longer respect the ‘upstream then downstream’ ordering rule).

### 2GS Datasets

The LOGAN approach was applied to the 2GS datasets, and created lossless, high efficiency representation thereof. Extracting the sequences after loading the LOGAN graph produced sequences identical to the input sequences, confirming the lossless property.

The construction times and resulting graph sizes are shown in Table 2.

**Table 2:** LOGAN graphs from 2GS: A LOGAN graph was created from each 2GS dataset. Build time was measured using a machine equipped with a 4-core, 3.5GHz (3.9GHz turbo) E3-1240V5 processor.

| Dataset | Reads (m) | Bases (Mbp) | Build Time (s) | Graph Size (MB) |
| --- | --- | --- | --- | --- |
| *E. coli* 1 | 7.6 | 543.7 | 13.6 | 101.0 |
| *E. coli* 2 | 221.0 | 19,743.8 | 1394.0 | 617.9 |
| *A.thaliana* | 30.4 | 2,364.5 | 133.5 | 478.3 |
| *S. pennellii* | 450.3 | 56,657.0 | 9,319.2 | 6,345.8 |

### 3GS and Hybrid Datasets

The LOGAN approach was also verified on a 3GS dataset alone, and as a hybrid combination with a 2GS dataset. The construction times and resulting graph sizes are shown in Table 3.

**Table 3:** LOGAN graphs from 3GS and hybrid datasets: A LOGAN graph was created from the *E. coli* 3GS dataset alone, and in combination with an *E. coli* 2GS dataset.

| Dataset | Reads (m) | Bases (Mbp) | Build Time (s) | Graph Size (MB) |
| --- | --- | --- | --- | --- |
| *E. coli* PB1 | 0.15 | 1,060.5 | 214.1 | 1,290.7 |
| *E. coli* 1 & PB1 | 7.6 & 0.15 | 543.7 & 1,060.5 | 232.5 | 1,380.4 |

## Discussion

The LOGAN graph promises to be the foundation of a novel approach to large scale sequence analysis, by providing a lossless in-memory representation of the dataset, suitable for *de novo* assembly or other analysis. It avoids the main limitations of existing approaches, and in principle allows flexible integration of 2GS, 3GS and pre-assembled data including reference data. Unlike the majority of reference-based approaches, the LOGAN approach works equally well with multiple or partial references, and does not introduce bias if used with reference sequences from relatively distant relatives, such as those from another species. The lossless nature of the LOGAN graph also lends itself well to datasets with high dynamic range, including those from transcriptomic and meta-genomic projects, since no early coverage-based filtering is needed.

The current implementation can create lossless sequence graphs from 2GS datasets of gigabase-scale genomes in an acceptable time on a modest desktop or workstation. Compared to the DBG approach, the lossless nature of the LOGAN graph enables more efficient usage of the information in a dataset, especially for challenging datasets. This avoids the various trade-offs inherent in the selection of *k*, the node size, while also avoiding the need to iteratively build multiple graph representations, such as in the SPAdes ^(21)^ multi-kmer approach. Furthermore, despite its greater information content, the representation provides dramatically better memory efficiency than many established DBG-based assemblers, and is broadly competitive in memory terms with state of the art approaches such as Succinct Graphs ^(11)^ and Minia ^(13)^. Finally, unlike the DBG approach, the error tolerant and lossless nature of the LOGAN graph makes integration of 3GS data feasible.

Since the LOGAN graph is relatively tolerant of sequence errors, 3GS and hybrid datasets are also supported. This enables the combined single-stage analysis of hybrid datasets, avoiding the information loss inherent in multi-step approaches. That said, the current implementation is primarily tuned to the characteristics of 2GS datasets. Future modifications, including sparser indexing with longer edges and more refined selection of s-mer nodes should enable dramatic improvements in 3GS representational efficiency.

Unlike OLC-based approaches, the LOGAN graph does not enforce a particular sequence of analysis steps, and thus is suitable for a data-centric, iterative refinement approach. The use of a single representation throughout the analysis avoids much of the computational overhead inherent in the hierarchical or iterative application of the OLC method, while the in-memory sequence representation dramatically reduces the need to access external memory.

Building the LOGAN graph is, of course, only the first step towards an improved sequence analysis approach, and later steps such as error correction, clustering and path finding will be required to form a complete *de novo* assembly solution. However, unlike existing approaches, the LOGAN graph does not limit the later analysis steps to a lossy view or subset of the data. This greater informational access should translate to improved results, especially with more challenging datasets and complex targets. This potential has already been validated by using an earlier implementation of the LOGAN graph approach as the basis of prototype error corrector for RNA-Seq data, PAGANtec ^(22)^. This was shown to outperform existing approaches, despite inefficiencies in the preliminary graph implementation.

## Conclusion

The LOGAN graph provides the first step towards a ‘grand unified’ sequence analysis approach by enabling the lossless in-memory representation of large-scale datasets from 2GS and/or 3GS platforms, optionally supplemented by reference or other pre-assembled data. The existing implementation shows high representational efficiency and can process realistic 2GS datasets from gigabase-scale genomes such as *S. pennellii* in hours on a desktop machine. The representational efficiency and runtime can be expected to improve further as additional optimizations are implemented. It has also been validated on 3GS and hybrid datasets, but further 3GS specific tuning is needed to improve the representational efficiency and build time with these datasets.

## References

[1] Paul Medvedev and Michael Brudno. Maximum likelihood genome assembly. Journal of computational Biology, 16(8):1101–1116, 2009.

[2] Saul B Needleman and Christian D Wunsch. A general method applicable to the search for similarities in the amino acid sequence of two proteins. Journal of molecular biology, 48(3): 443–453, 1970.

[3] Temple F Smith and Michael S Waterman. Identification of common molecular subsequences. Journal of molecular biology, 147(1):195–197, 1981.

[4] Jason R Miller, Sergey Koren, and Granger Sutton. Assembly algorithms for next-generation sequencing data. Genomics, 95(6):315–327, 2010.

[5] Sergey Koren and Adam M Phillippy. One chromosome, one contig: complete microbial genomes from long-read sequencing and assembly. Current opinion in microbiology, 23:110–20, 2015.

[6] Eugene W Myers. The fragment assembly string graph. Bioinformatics, 21(Suppl 2):ii79–ii85, 2005.

[7] Michael Burrows and David J Wheeler. A block-sorting lossless data compression algorithm. 1994.

[8] Jared T Simpson and Richard Durbin. Efficient de novo assembly of large genomes using compressed data structures. Genome research, 22(3):549–556, 2012.

[9] Sergey Koren, Brian P Walenz, Konstantin Berlin, Jason R Miller, Nicholas H Bergman, and Adam M Phillippy. Canu: scalable and accurate long-read assembly via adaptive k-mer weighting and repeat separation. bioRxiv, page 071282, 2017.

[10] Pavel A Pevzner, Haixu Tang, and Michael S Waterman. An eulerian path approach to dna fragment assembly. Proceedings of the National Academy of Sciences, 98(17):9748–9753, 2001.

[11] Thomas C Conway and Andrew J Bromage. Succinct data structures for assembling large genomes. Bioinformatics, 27(4):479–486, 2011.

[12] Guy Joseph Jacobson. Succinct static data structures. 1988.

[13] Kamil Salikhov, Gustavo Sacomoto, and Gregory Kucherov. Using cascading bloom filters to improve the memory usage for de brujin graphs. Algorithms for Molecular Biology, 9(1):2, 2014.

[14] Chengxi Ye, Zhanshan Sam Ma, Charles H Cannon, Mihai Pop, and W Yu Douglas. Exploiting sparseness in de novo genome assembly. BMC bioinformatics, 13(6):S1, 2012.

[15] Lee D Erman, Frederick Hayes-Roth, Victor R Lesser, and D Raj Reddy. The hearsay-ii speech-understanding system: Integrating knowledge to resolve uncertainty. ACM Computing Surveys (CSUR), 12(2):213–253, 1980.

[16] Paris Avgeriou and Uwe Zdun. Architectural patterns revisited–a pattern. In European Conference on Pattern Languages of Programs, 2005.

[17] Martin De Prycker. Asynchronous transfer mode: solution for broadband ISDN. Ellis Horwood, 1991.

[18] Eric Rosen, Arun Viswanathan, and Ross Callon. Multiprotocol label switching architecture. Technical report, 2000.

[19] Tokuhiro Tsukiyama, Yoshie Kondo, Katsuharu Kakuse, Shinpei Saba, Syoji Ozaki, and Kunihiro Itoh. Method and system for data compression and restoration, April 29 1986. US Patent 4,586,027.

[20] Anthony M Bolger, Marc Lohse, and Bjoern Usadel. Trimmomatic: a flexible trimmer for illumina sequence data. Bioinformatics, page btu170, 2014.

[21] Anton Bankevich, Sergey Nurk, Dmitry Antipov, Alexey A Gurevich, Mikhail Dvorkin, Alexander S Kulikov, Valery M Lesin, Sergey I Nikolenko, Son Pham, Andrey D Prjibelski, et al. Spades: a new genome assembly algorithm and its applications to single-cell sequencing. Journal of computational biology, 19(5):455–477, 2012.

[22] Markus Joppich, Dirk Schmidl, Anthony M Bolger, Torsten Kuhlen, and Björn Usadel. Pagan-tec: Openmp parallel error correction for next-generation sequencing data. In International Workshop on OpenMP, pages 3–17. Springer, 2015.

